# Genomic foundation models do not yet boost common variant discovery beyond standard functional annotations

**DOI:** 10.64898/2026.10.13.758880

**Authors:** Andrew J. Bass, Joshua Park, Sach Mukherjee, John Whittaker, Chris Wallace

## Abstract

Genomic foundation models (GFMs) use large-scale AI to model genetic sequences and their influence on molecular phenotypes. While GFMs have shown promise in rare variant analysis, their utility for common variant discovery remains untested. Using the surrogate functional false discovery rate framework for functionally informed GWAS discovery, we compared standard functional annotations (baselineLD) with annotations derived from three leading GFMs: the sequence-to-function models AlphaGenome and Borzoi, and the genomic language model GPN-Star. Across 27 UK Biobank traits, GPN-Star decreased locus discovery relative to a standard GWAS, whereas AlphaGenome and Borzoi increased discovery but by less than baselineLD (medians 1.8%, 3.8%, and 5.1%). When we combined the GFM and baselineLD annotations, there was no increase over baselineLD alone (median changes −0.1% to 0.2%), suggesting that AlphaGenome and Borzoi mostly capture information already in baselineLD. Thus, GFMs may require larger corpora or trait-specific feature engineering to improve discovery beyond standard functional annotations.

---

AI models are having a major impact on biomedical and health research. In particular, large language models (LLMs), other foundation models and multimodal variants are being applied to problems ranging from protein structure prediction to decoding the non-coding genome.^1,2^ For example, AlphaFold can accurately predict a protein’s 3D structure from its amino acid sequence,^3^ potentially saving months or years of experimental effort. While there are many models that allow us to accurately predict the consequences of protein-coding variation,^4^ the majority of trait-associated variants in genome-wide association studies (GWAS) are non-coding. As such, there is now considerable effort to apply deep learning models to the more difficult task of interpreting non-coding variation.

Characterizing non-coding variation is challenging because effects are indirect and context dependent, and describing them typically requires multimodal measurements. One well-established genome annotation is evolutionary constraint, which reflects how strongly natural selection acts to remove deleterious mutations, with at least 10.7% of the human genome estimated to be evolutionarily constrained across mammals.^5^ Similar whole-genome alignment data used to estimate constraint have been used to train LLM-like models (“genomic language models”; e.g., GPN-Star), which have been shown to identify functional elements such as enhancers.^6^ Moving beyond classical measures of evolutionary constraint, the goal of these and more recent genomic foundation models (GFMs) is to leverage large-scale training on sequence (and other biological) data to implicitly learn how sequence affects molecular phenotypes. Here, we use the term GFM broadly to include both self-supervised genomic language models (GPN-Star) and supervised sequence-to-function models (AlphaGenome and Borzoi). For example, Borzoi is trained to predict multimodal functional data (e.g., RNA-seq and DNA accessibility) at a 32 base-pair (bp) resolution,^7^ while AlphaGenome provides similar predictions at up to single bp resolution.^8^ Variant scores derived from Borzoi have already been integrated into fine mapping of common disease GWAS signals, demonstrating their utility for complex trait genetics.^9^ These results align with work that has shown that incorporating experimentally derived functional data, a proxy for a variant’s downstream impact, improves rare variant analysis and functional fine mapping.^10,11^

While GFMs have already proven capable of predicting molecular features of the genome, their utility for downstream genetic applications remains an active area of research. Since the vast majority of trait-associated variants map to non-coding regions, GFMs have the potential to provide a comprehensive set of variant effect predictions, computationally derived scores capturing the functional relevance of genetic variation. However, the extent to which the representations obtained by GFMs capture information beyond existing approaches to functional annotation remains an open question, with earlier work showing deep learning models provided limited information for disease association.^12^ Such observations suggest that the utility of GFMs for specific tasks requires rigorous and targeted evaluation and cannot simply be assumed from accuracy in predicting underlying regulatory annotations. Given that standard functional annotations can substantially increase power in GWAS,^13^ we assess whether current GFM scores can also provide information beyond these annotations to improve common variant discovery. To our knowledge, such an assessment has not been previously carried out for GFMs, and differs from prior benchmarks which focused on heritability enrichment,^12^ rare variant testing to evaluate coding variants, and fine mapping to prioritize variants among a set of associated variants. Common variant discovery is potentially a more difficult task that requires predicting the effect of non-coding variation with sufficient accuracy to prioritize sub-genome-wide significant signals without increasing the type I error rate.

We use the recently developed surrogate functional false discovery rate (sfFDR) framework^14^ for functionally informed GWAS discovery. sfFDR has been shown to improve power by incorporating auxiliary information, such as GWAS summary statistics for a related trait, while controlling the type I error rate. sfFDR is flexible in the sources of information it can incorporate, making it a natural approach to integrate the output of diverse GFMs within a principled statistical framework. Specifically, we assess the utility of GFM-derived features for GWAS discovery relative to the baselineLD annotations.^15^ We hypothesize that if GFM scores contain *additional* information, then incorporating them should improve GWAS discovery.

We assessed whether GFM predictions improve discovery compared with standard annotations (Table 1) by analyzing 27 approximately independent traits from the UK Biobank^13^ covering four general categories, namely, anthropometric, blood cell, disease, and other (Methods). Across the traits, the proportion of SNP heritability explained by lead SNPs was modest (median 38.1%), with disease traits capturing the lowest (26.5%) and blood cell traits the highest (67.8%; Figure 1A). The gap in lead SNP heritability suggests an opportunity to improve discovery of association signals by leveraging functional data.

**Figure 1:**
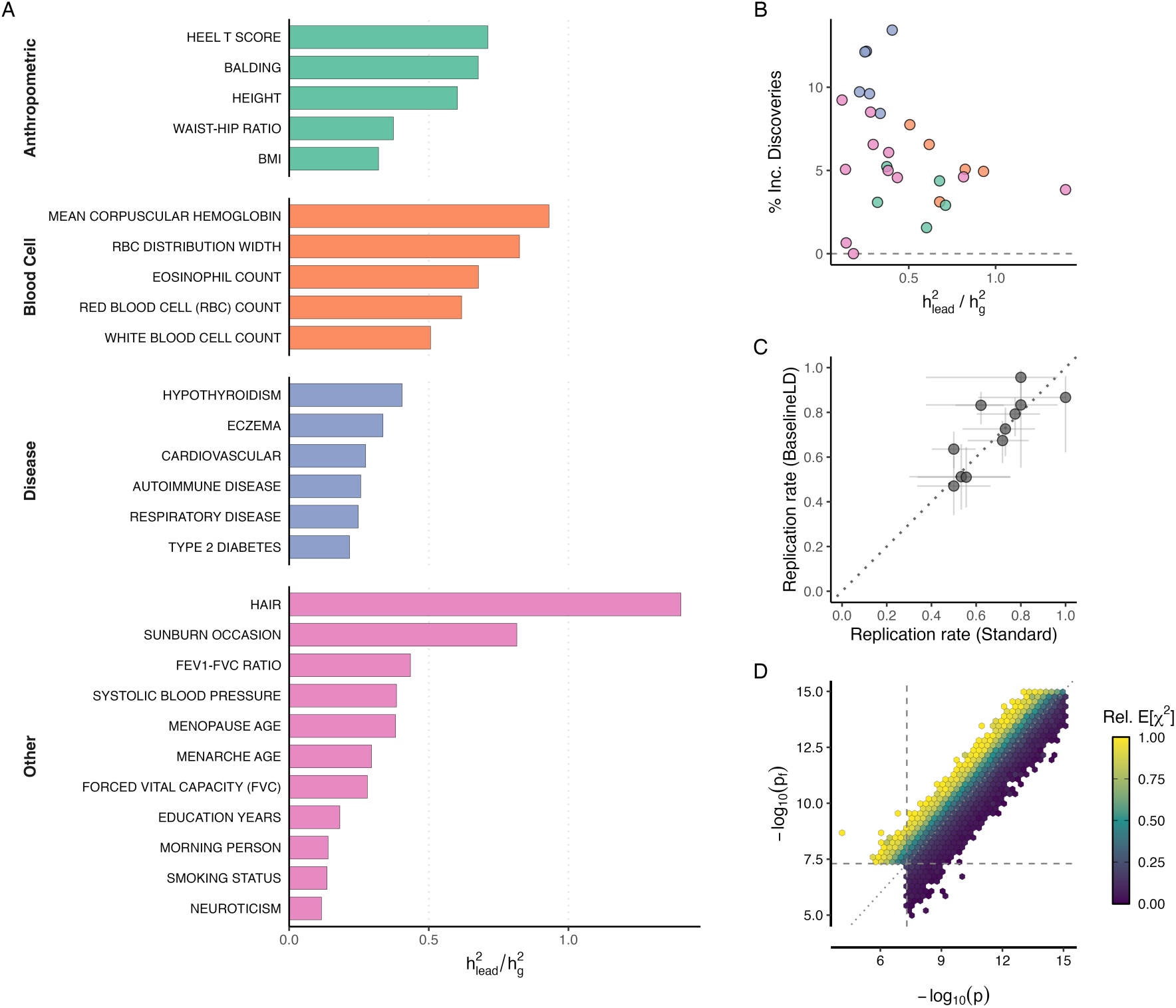
Performance and validation of the sfFDR framework across 27 approximately independent UK Biobank traits. (A) Proportion of SNP heritability explained by lead variants 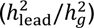 across multiple trait categories (denoted by color). (B) Relative increase in the discovery of independent loci using functional *p*-values compared with standard *p*-values as a function of 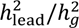. (C) The replication rate (*p* < 0.05) of the distinct loci identified by only the functional *p*-values (y-axis) compared with those identified only using the standard *p*-values (x-axis). Error bars denote the 95% confidence intervals. (D) Relationship between significant functional and standard *p*-values across all traits. The hexagons are colored by the mean expected *χ*^2^ predicted from the baselineLD annotations (scaled within each trait), which are used in the sfFDR framework to calculate functional *p*-values. Note that any values of −log_10_(*p*) larger than 15 are shown at 15.

**Table 1:**
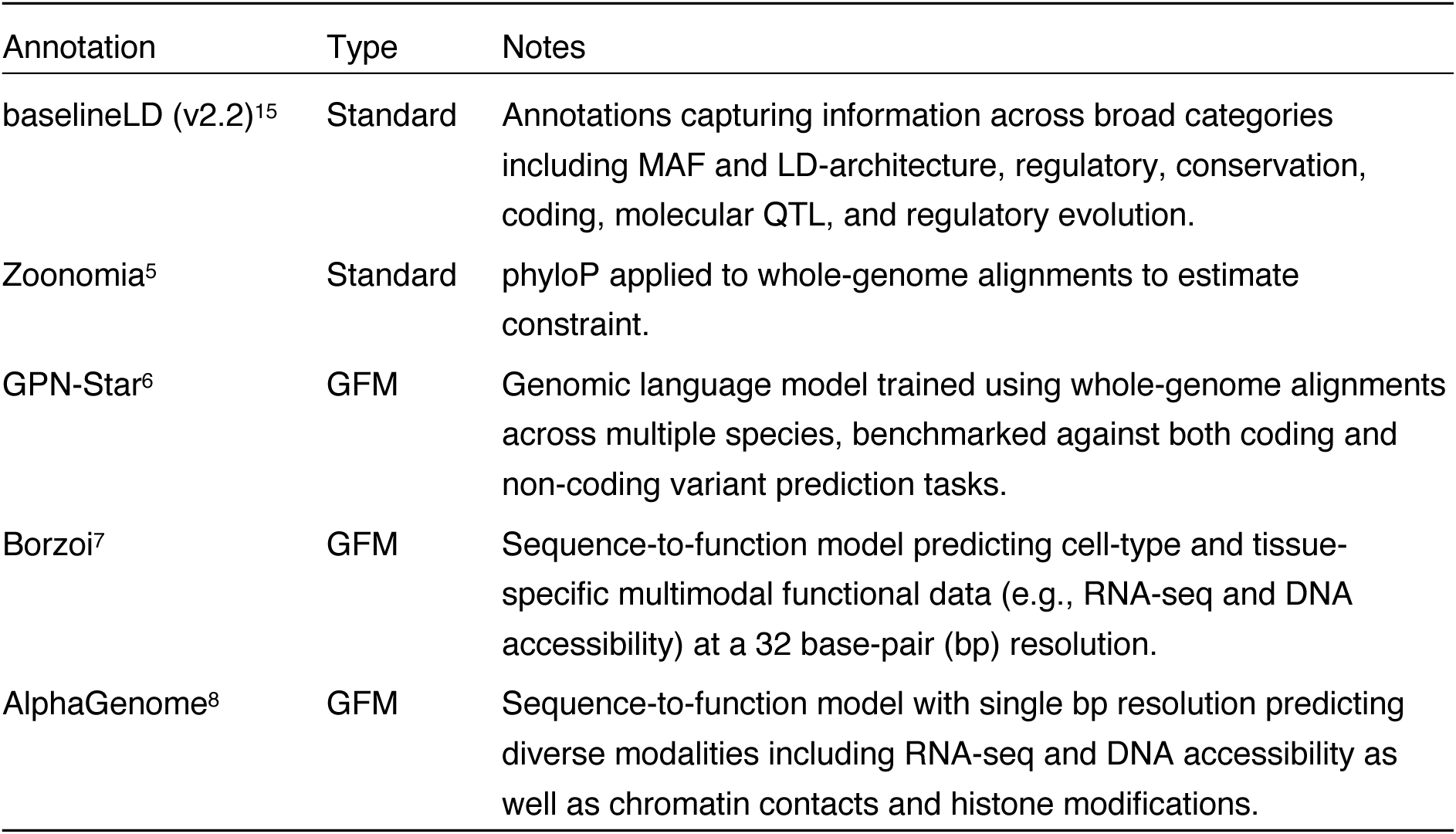
Annotations compared for their ability to enhance GWAS discovery. GFM, genomic foundation model.

| Annotation | Type | Notes |
| --- | --- | --- |
| baselineLD (v2.2)<sup>15</sup> | Standard | Annotations capturing information across broad categories including MAF and LD-architecture, regulatory, conservation, coding, molecular QTL, and regulatory evolution. |
| Zoonomia<sup>5</sup> | Standard | phyloP applied to whole-genome alignments to estimate constraint. |
| GPN-Star<sup>6</sup> | GFM | Genomic language model trained using whole-genome alignments across multiple species, benchmarked against both coding and non-coding variant prediction tasks. |
| Borzoi<sup>7</sup> | GFM | Sequence-to-function model predicting cell-type and tissue-specific multimodal functional data (e.g., RNA-seq and DNA accessibility) at a 32 base-pair (bp) resolution. |
| AlphaGenome<sup>8</sup> | GFM | Sequence-to-function model with single bp resolution predicting diverse modalities including RNA-seq and DNA accessibility as well as chromatin contacts and histone modifications. |

To incorporate annotation data into the sfFDR framework, we first summarized the data into a single expected chi-squared variable, following a similar procedure to stratified LD score regression.^16^ We considered three metrics to evaluate power improvements. First, we assessed the ability of the GFM-derived metric to predict the strength of SNP association: the out-of-sample (OOS) *R*^2^ of the ridge regression model used to generate the expected chi-squared values. Second, since sfFDR uses the chi-squared variable to inform the ranking of SNPs, we calculated the signal enrichment: the mean observed chi-squared of the top 1%, 5%, and 10% of SNPs ranked by each model relative to baselineLD. Third, we conservatively defined discoveries as genome-wide significant (*p* < 5 × 10^−8^) and uncorrelated (*r*^2^ < 0.01) lead SNPs. Thus, we evaluated each annotation set on whether it improves prediction of GWAS association, how influential the predicted chi-squared is in reranking SNPs, and whether that translates into additional genome-wide significant loci.

Across traits, incorporating baselineLD increased the number of discoveries (median increase 5.07%; Figure 1B). In particular, the increase was largest for disease traits (median 10.9%), which had the smallest proportion of heritability captured by lead SNPs. Importantly, the distinct discoveries identified by sfFDR replicated in independent data from FinnGen (available for 11 of the 27 traits) at a comparable rate to those only discovered by the standard analysis (Figure 1C, Table S1). Among the 11 traits, sfFDR identified 657 distinct discoveries (not found by standard analysis) replicating at a rate of 0.69 (95% CI 0.65–0.72) compared with 347 distinct discoveries from standard *p*-values (0.61, 95% CI 0.56–0.66). The 6,673 loci identified by both approaches replicated at 0.84 (95% CI 0.83–0.85). These additional discoveries arise because sfFDR alters the ranking of individual SNPs by prioritizing SNPs with high expected chi-squared values and deprioritizing those with low values (Figure 1D).

The increase in discoveries was smaller with Borzoi and AlphaGenome on their own than with baselineLD (median increase 3.80% and 1.78%, respectively, compared with 5.07%), while the number decreased with Zoonomia and GPN-Star (Figure 2A). We found that all annotations produced positive out-of-sample *R*^2^ values that were mostly lower than baselineLD’s (Figure 2B). The constraint annotations were significant in all of the traits but explained the least variance, and the SNPs they ranked highest were similarly or less strongly associated (median observed chi-squared 5.8 for Zoonomia and 4.2 for GPN-Star in the top 1%, compared with 5.7 for AlphaGenome, 6.8 for Borzoi and 8.0 for baselineLD; Figure 2C, D).

**Figure 2:**
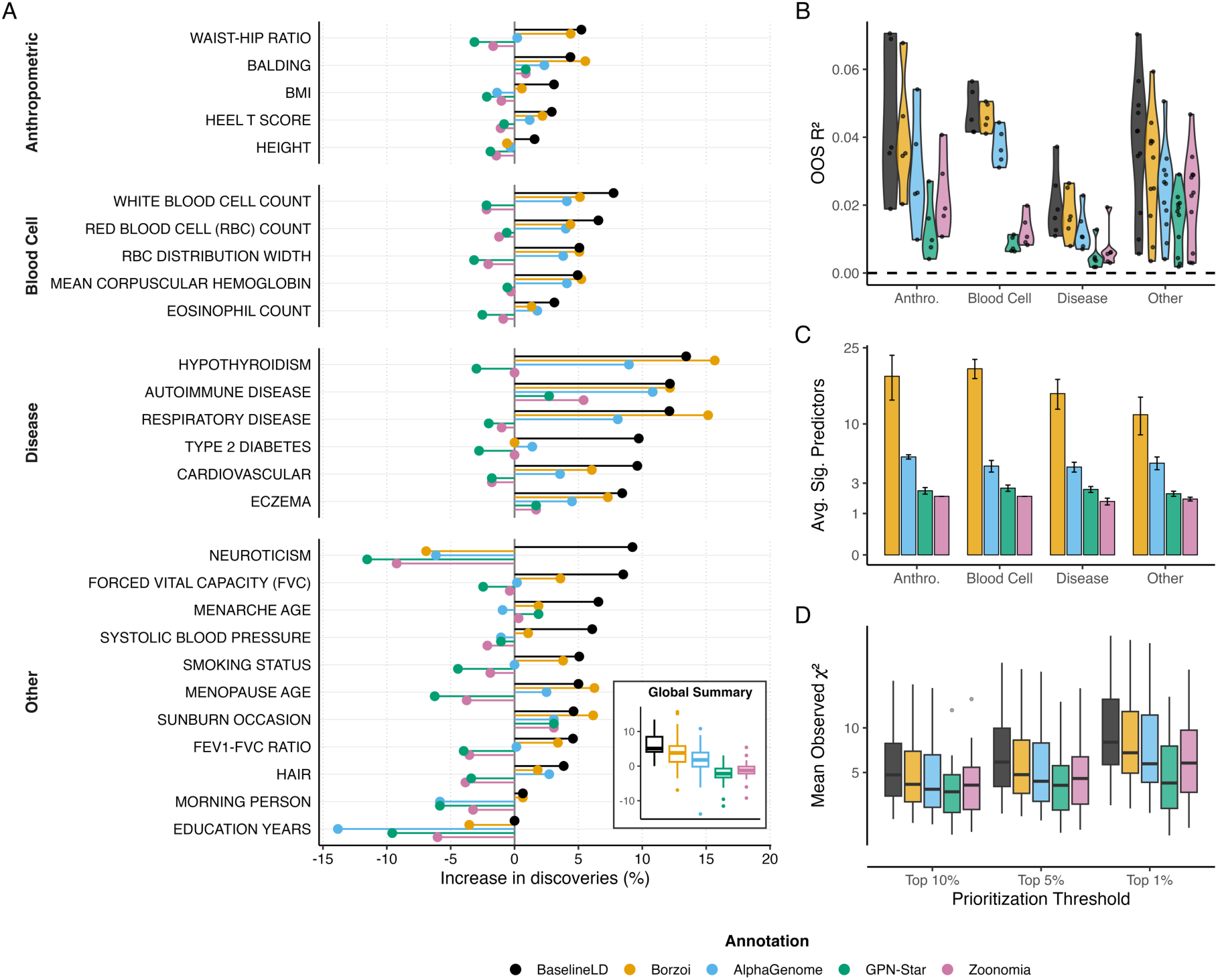
Impact of annotation sets used on their own on common variant discovery. (A) Relative increase in the discovery of independent loci using functional *p*-values compared with standard *p*-values across the annotation models (color). (B) A violin plot of the out-of-sample (OOS) *R*^2^ for each annotation model by trait class. Points denote individual traits. (C) The average number of significant explanatory variables (FDR < 0.2) across trait classes (square root scale). Note that there were 2, 3, 8, and 102 annotations from Zoonomia, GPN-Star, AlphaGenome, and Borzoi, respectively. Error bars show ±1 s.e.m. across traits. (D) Boxplots of the average observed *χ*^2^ across the top 1%, 5%, and 10% of variants prioritized by each model (pseudo log scale). Points denote traits that are outliers.

Borzoi annotations have previously been used in combination with the baselineLF annotations (an extension of baselineLD to low frequency variants).^9^ We found that adding each annotation set to baselineLD performed similarly to using baselineLD alone (Figure 3A). The median difference in lead SNPs across traits between the combined annotation and baselineLD alone ranged from −0.1% (AlphaGenome) to 0.2% (Borzoi). In total across the 27 traits, the combined annotations identified 13,416 to 13,447 loci compared with 13,420 for baselineLD alone. Despite only marginal changes in discovery, the combined annotations did improve OOS *R*^2^ compared with baselineLD alone (Figure 3B), and each model contributed at least one of its own annotations significant at a false discovery rate (FDR) < 0.2 in 15 to 24 of the 27 traits (Figure S1). In particular, Borzoi substantially increased the OOS *R*^2^ across all traits (average relative increase of 11.2%; Figure 3B). The improvements varied across traits and were not concentrated in traits where baselineLD was least predictive (Figure 3C). Signal enrichment was also higher in the baselineLD + Borzoi model than in baselineLD alone (Figure 3D).

**Figure 3:**
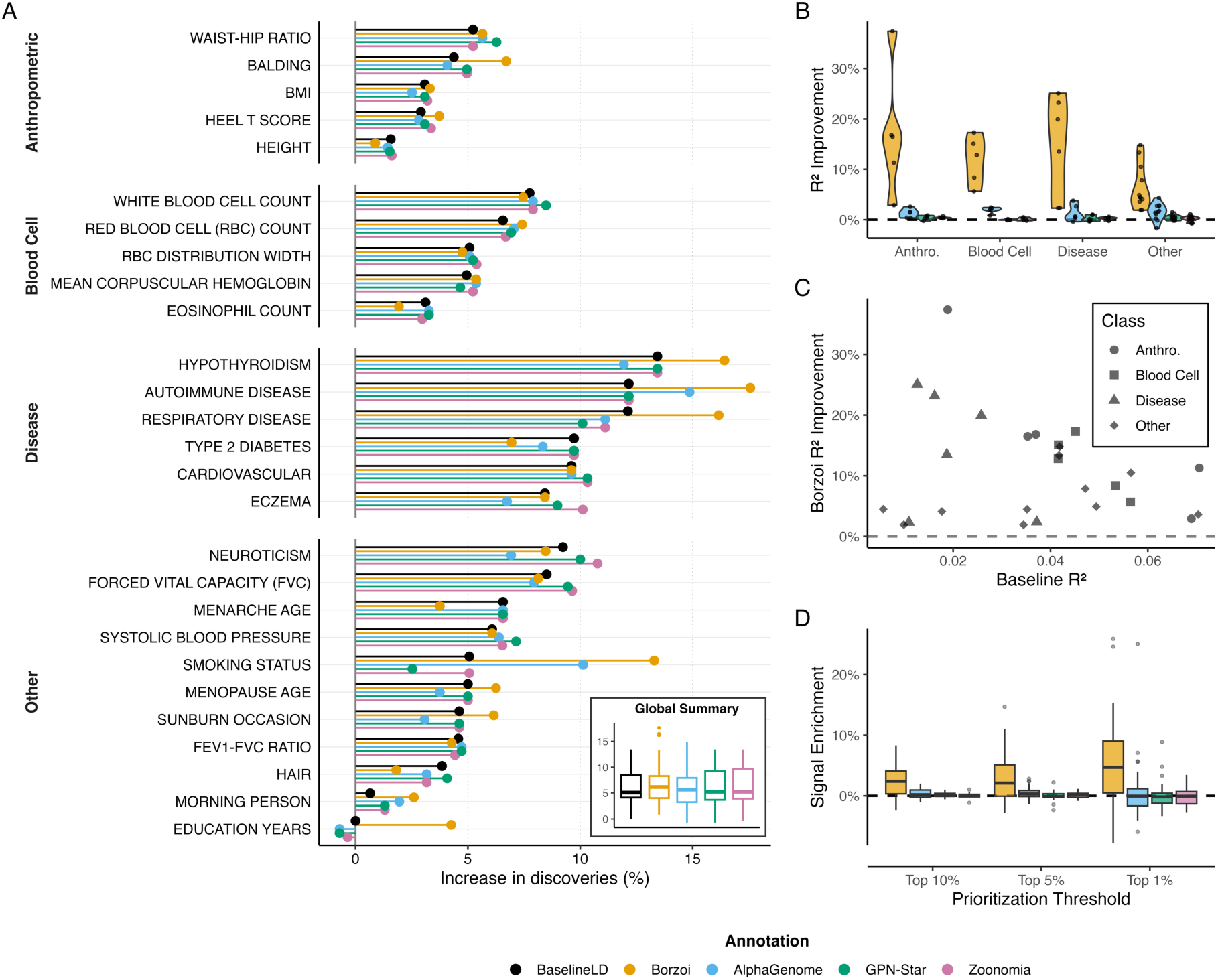
Evaluating foundation model annotations in combination with baselineLD. (A) Relative increase in the discovery of independent loci using functional *p*-values compared with standard *p*-values. Note that baselineLD annotations were added to each model annotation (color). (B) Relative out-of-sample (OOS) *R*^2^ improvement over baselineLD by trait class. (C) Borzoi relative OOS *R*^2^ improvement compared with baselineLD OOS *R*^2^. Trait class is denoted by shape. (D) Signal enrichment across annotation models. Signal enrichment is the relative improvement over baselineLD in the average observed *χ*^2^ across the top 1%, 5%, and 10% of variants prioritized by each model.

We next evaluated the overlap between Borzoi and baselineLD annotations. We found that Borzoi’s assay categories were strongly correlated with the baselineLD regulatory annotations (Figures 4A and S2A). However, after conditioning on the regulatory category, the variance explained by, and partial correlations with, the other baselineLD categories (conservation, coding, molecular QTL, regulatory evolution, and MAF and LD-architecture) were minimal (Figures 4B and S2B). In particular, the partial correlations with evolutionary constraint were close to zero, even though a constraint annotation was the most frequently significant functional annotation in the ridge regression model used to estimate the chi-squared variable in sfFDR (significant in 59% of traits; the most frequently significant annotations overall were MAF and LD-architecture terms; Figure S1). Overall, these results suggest that Borzoi’s contribution substantially overlaps with the regulatory information in baselineLD.

**Figure 4:**
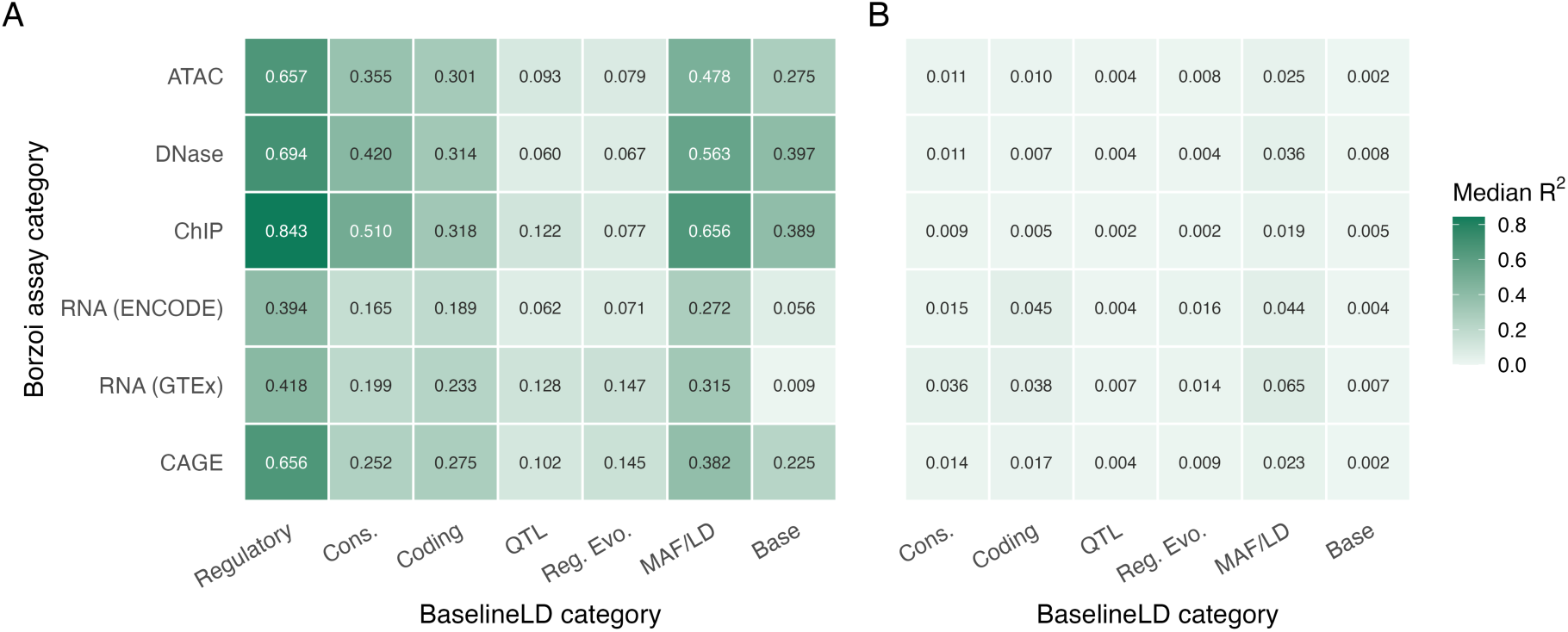
Borzoi overlaps substantially with baselineLD regulatory annotations. Median variance explained (*R*^2^) in the LD scores of Borzoi principal components (PCs) by baselineLD annotation categories. The rows are Borzoi’s six assay categories and the columns are baselineLD annotation categories (Base denotes the total LD score). For each PC, *R*^2^ was obtained by regressing its LD score on the LD scores of all annotations within a category. The tiles show the median across the principal components of that assay category. (A) The marginal *R*^2^ of each category. (B) The added variance explained after conditioning on the regulatory baselineLD annotations.

We therefore tested whether discovery is limited by the regulatory annotation class itself or by Borzoi’s representation of it. We applied sfFDR to nested annotation subsets that share the same 17 baselineLD MAF and LD-architecture annotations, i.e., annotations that do not contain functional information (Methods). Across all traits, the MAF and LD-architecture annotations identified 318 additional loci, which is roughly half of the baselineLD model (598). When we added one functional class at a time, we found that the regulatory annotations from baselineLD, the AlphaGenome and Borzoi scores, the Zoonomia and GPN-Star constraint scores, and the 10 conservation annotations from baselineLD all increased discovery (Figure S3). In particular, adding Borzoi’s annotations identified 545 additional loci (56 more than baselineLD’s regulatory annotations). Our subset analysis showed that regulatory annotations from Borzoi provided the largest increase in loci of any annotation class when added to the MAF and LD-architecture annotations, indicating that predicted molecular phenotypes are informative for common variant discovery. However, this combination did not improve power over the full baselineLD model (median increase 5.08% compared with 5.07%; Figure S3).

Across our analyses, increases in signal enrichment and OOS *R*^2^ did not always translate into more discoveries. We believe this reflects the heterogeneous genetic architecture of common traits: some variants will lie in constrained regions, some variants will act through changing proteins, and some variants will act through regulatory mechanisms. Against this patchwork, a single class of annotations can detect real enrichment which can improve prediction and increase OOS *R*^2^. However, the reranking performed by sfFDR produces both winners and losers: when the annotations are biased to a single class (e.g., constraint with Zoonomia or regulation with Borzoi) then the SNPs in that class are pushed towards significance (the “winners”) but may be offset by SNPs outside the class that are pushed away. Since baselineLD spans multiple classes, it is less affected by this trade-off.

While results with baselineLD confirm that molecular phenotypes do inform common variant discovery, and while GFMs may capture certain aspects of function or constraint, they did not add to standard functional annotations in our benchmark. We assume that there is a path from molecular readouts such as transcription and chromatin accessibility, to whole-organism phenotypes, to the tolerance of individual bases to mutation across evolution, but these are complex relationships, involving functional genomic outputs and their interplay with environmental factors across a lifetime or across generations. Hence, GFMs may require much larger data corpora than current training sets (both more genomes and more functional readouts) to fully annotate genome features.

Alternatively, while GFM papers often report results with zero-shot predictions, additional fine-tuning or feature engineering steps are common. Here, to best reflect the use of GFMs in an off-the-shelf manner for a downstream task, we focused on integrating GFM output without any optimization of the GFMs themselves. Our results suggest that additional work will be needed to make the most of GFMs in this setting, for example feature engineering, possibly in a trait (class)-specific manner to extract information from these models for GWAS discovery. There are several options that could be explored. Distillation is the process of compressing the high-dimensional, often highly correlated output tracks of a foundation model into a smaller, denser set of features to reduce statistical noise and isolate the core biological signal. Borzoi scores were distilled through principal component analysis (PCA) by the Sniff pipeline, while for AlphaGenome we took the maximum absolute score within each output type. Thus, the comparison between the two models reflects the representations as much as the models themselves. Given the performance difference, it would be interesting to explore whether PCA or other options such as variational autoencoders may capture more information from AlphaGenome. It is also possible that restricting the tissue-specific scores to trait-relevant tissues before distillation could further improve performance.

Fine-tuning is the process of adapting a pre-trained model’s weights (and thereby generalized scores) to optimize them for a specific downstream analytical objective. Fine-tuning has been shown to improve performance of deep learning model predictions to prioritize rare non-coding variants^17^ and to improve the prediction of gene expression at the individual level by GFMs.^18^ However, the training objective matters: the recently released AlphaGenome variant impact score (AVI),^19^ which combines AlphaGenome and AlphaMissense predictions with conservation and other scores to separate rare from common variants, did not improve discovery when added to baselineLD (median increase 5.23% compared with 5.07%, and 13,422 and 13,420 loci in total; Figure S3). GFM scores could instead be fine-tuned for common variant discovery, for example through training a lightweight model to predict GWAS expected chi-squared statistics from the GFM features, or through established efficient fine-tuning approaches such as low-rank adaptation.^20^

Despite the distance from functional genome outputs to human phenotypes, learning how to optimally use GFMs may help with many post-GWAS tasks, and we suggest that optimizing the distillation and fine-tuning of tracks for enhancing common variant discovery using approaches such as sfFDR may inform how these tracks should best be used for those tasks. For example, GFMs could improve polygenic risk prediction, where there is a recognized and persistent bias in existing GWAS resources towards European ancestry,^21^ which translates into weaker performance in individuals of non-European ancestries.^22^ While GWAS require large sample sizes, GFMs require a diversity of tracks, each with relatively few examples. Thus, careful selection of tracks when GFMs are trained may provide ancestry-agnostic predictions of function from sequence, which can then be incorporated into polygenic risk scores to counter European bias.

We conclude that, for the task of increasing common variant GWAS discovery, GFMs do not yet outperform standard annotations of function or constraint. Nevertheless, GFMs represent a major advance in computational models for genome analysis and are likely to be critical components of many future analyses. To inform their effective use in genomics, their utility must be evaluated for each downstream task. Going beyond common variant discovery, we recommend that AI foundation models in general, and GFMs in particular, be tested in a detailed, domain-specific fashion using carefully constructed metrics, and compared with existing approaches to understand their added value. Such research is essential both for the domain-specific adoption of AI models and for informing their further development.

## Methods

### sfFDR framework extended to functional annotations

#### Overview

We integrated the GFM annotations using the recently developed surrogate functional false discovery rate (sfFDR) framework,^14^ implemented in the sffdr software (v1.1.2). The sfFDR framework provides estimates of the functional local false discovery rate and *q*-value,^23^ and uses these quantities to construct a functional *p*-value to control the type I error rate. Functional *p*-values integrate information from auxiliary data while testing the same null hypothesis as the original GWAS, and can therefore be interpreted in the same manner as standard *p*-values. While the sfFDR framework was originally formulated to incorporate summary-level data for genetically related traits (i.e., pleiotropy), the methodology and underlying theory extend to handle any type of informative data. Here, we extended the procedure to incorporate SNP-level functional annotations.

We briefly review the extension of sfFDR to functional data. Consider a set of *p*-values, *p_i_*, and a set of informative functional data, ***z****_i_*, for *i* = 1,2, …, *m* SNPs in a GWAS. For each SNP, sfFDR estimates the functional local FDR, **Λ**(*p*, ***z***), which is the probability that a hypothesis is null given its observed *p*-value and associated informative data:

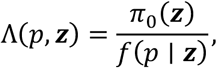

where *π*_0_(***z***) is the conditional prior probability of a SNP being null for the trait of interest and *f*(*p* ∣ ***z***) is the conditional density of the *p*-values given the informative data. We can then define a decision rule to determine a subset of significant SNPs based on the functional local FDR and a significance threshold *τ* as Γ*_τ_* = {(*p*, ***z***): **Λ**(*p*, ***z***) ≤ *τ*}. Previous work has shown that this decision rule is the Bayes rule minimizing a *τ*-weighted combination of the probabilities of a false positive and a false negative.^23^

Based on the above decision rule, we can map these quantities to a functional *p*-value to control the type I error rate:

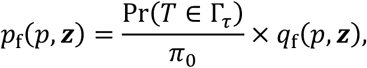

where *τ* = **Λ**(*p*, ***z***) is the functional local FDR of the SNP being tested, *π*_0_ = E***_z_***C*π*_0_(***z***)D is the prior probability of a null SNP, Pr(*T* ∈ Γ*_τ_*) is the probability of the rejection region under the distribution of *T* = (*p*, ***z***), and 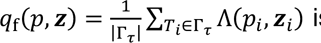 is the functional *q*-value. The functional *p*-value can also be written as 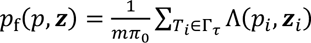, where *π*_0_ is estimated by 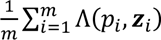, and so the *q*_f_-value normalizes by the number of rejections while the *p*_f_-value normalizes by the total null mass. Thus, the functional *p*-value depends on the functional local FDR, which depends on the conditional prior probabilities and the conditional density. Notably, the functional *p*-value is a monotone transform of the functional local FDR, and so it preserves the Bayes-optimal ordering.

In sfFDR, the prior probabilities are estimated using a generalized linear model (GLM) and the conditional density is estimated non-parametrically using local likelihood kernel density estimation.^14^ A key feature of the sfFDR framework is that it reduces the dimensionality of this non-parametric density estimation by constructing a surrogate variable based on *π*_0_(***z***). Since the original work focuses on leveraging pleiotropy from GWAS summary statistics, we detail below how functional data are integrated to model the prior probabilities, while the corresponding estimation procedure remains identical to the original framework.

#### Modeling the prior probability *π*_0_(*z*) in sfFDR

To model the prior probabilities, we adapted the stratified LD score regression (S-LDSC) framework to estimate a SNP’s expected chi-squared statistic, E[*χ*^2^], conditional on functional data.^15,16^ This variable serves as a surrogate that summarizes the genetic contribution from the annotations, and has previously been used to reweight GWAS *p*-values.^13^ Specifically, for *k* = 1,2, …, *C* annotations, we constructed an LD annotation score 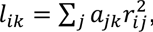, where *a*_j*k*_ is the *k*th annotation for SNP *j* and 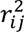 is the squared correlation between SNPs *i* and *j*. Intuitively, the LD annotation score incorporates local LD structure to account for signals captured by nearby SNPs. To compute the LD annotation scores, we used the PolyFun software^11^ with 489 European-ancestry individuals from the 1000 Genomes Project (Phase 3) as the reference panel.

Previous work^16^ has shown that the expected chi-squared statistic for the *i*th SNP can be modeled as

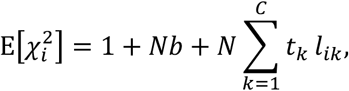

where *N* is the sample size, *t_k_* is the coefficient of the *k*th annotation, and *b* measures the contribution of confounding (e.g., due to population structure). We estimated these coefficients and the intercept (1 + *Nb*) using a weighted regression model on the HapMap3 SNPs (see ref. 16 for details). The MHC region was removed before model fitting, and SNPs with *χ*^2^ > max(0.001*N*, 80) were excluded from model fitting and evaluation. Note that each SNP’s weight depends on its LD score summed over all annotations. For annotation sets without baselineLD, this sum excluded the AlphaGenome, Borzoi, and AVI scores as their scores are standardized, and so their LD scores can be negative and destabilize the weights. To avoid instability caused by highly correlated annotations, we fitted a ridge regression model to predict expected chi-squared values (sets with fewer than five annotations were fitted by weighted least squares). Furthermore, to prevent data leakage within the sfFDR framework, we implemented a leave-one-chromosome-out (LOCO) cross-validation strategy: coefficients for a given chromosome were estimated using the remaining 21 chromosomes where each of the 21 training chromosomes was treated as a distinct fold when selecting the ridge penalty.

From the regression, we calculated the expected chi-squared, EO*χ*^2^P, for each SNP given the functional data. Since each chromosome was held out, we evaluated model performance using a weighted (i.e., using inverse LD score weights) out-of-sample (OOS) *R*^2^ on the HapMap3 SNPs. To assess whether the top ranked SNPs of each model were more strongly associated, we ranked the HapMap3 SNPs based on the estimated OOS expected chi-squared, and then calculated the weighted mean observed chi-squared of the top 1%, 5%, and 10% of ranked SNPs. We expressed this value relative to baselineLD to define the signal enrichment. The LOCO folds were also used as a block jackknife to calculate annotation standard errors 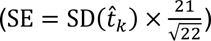 and corresponding *p*-values. We then assessed statistical significance across annotations within each trait at an FDR of 0.2.

We then estimated the prior probabilities in the sfFDR procedure using the expected chi-squared, where we flexibly modeled the rank-transformed expected chi-squared using a natural cubic spline basis to capture non-linearity. We placed knots at the rank of E[*χ*^2^] = 1 and at the 0.25, 0.5, 0.75, 0.9, 0.95, 0.99, and 0.999 quantiles of ranks above it, removing knots until each interval contained at least 100 LD-independent SNPs.

#### Functional annotations and variant effect predictions

We used the baselineLD (v2.2) model as our benchmark to evaluate the added value of GFMs. This dataset included 96 annotations for approximately 10 million variants from the 1000 Genomes Project (Phase 3) and corresponding reference files for 489 individuals of European ancestry.^15^ We also analyzed subsets of the baselineLD annotations: MAF and LD-architecture (MAF bins, MAF-adjusted predicted allele age, MAF-adjusted LLD-AFR, MAF-adjusted ASMC, recombination rate, nucleotide diversity, background selection statistic, and CpG content^15^), regulatory (histone marks, DHS, enhancers, promoters, transcription factor binding sites, and CTCF sites), conservation (GERP scores and constraint annotations across species), coding (coding, intronic, UTR regions, and synonymous and non-synonymous variants), molecular QTL (MaxCPP fine-mapping posteriors from GTEx and BLUEPRINT), and regulatory evolution (promoters, enhancers, and ancient sequence age). The six categories had 17, 44, 10, 10, 4, and 11 annotations, respectively.

We also considered scores from several recent models. This allowed us to quantify the new information gained relative to established annotations. We evaluated the following scores:

- **AlphaGenome**: For each variant, AlphaGenome (https://deepmind.google.com/science/alphagenome/) was queried using a 1 Mb sequence window centered on the variant. Variant effect scores were computed across eight output types spanning six functional modalities: RNA expression (RNA-seq), transcription start site activity (CAGE, PRO-cap), chromatin accessibility (ATAC-seq, DNase-seq), transcription factor binding (ChIP-TF), histone modifications (ChIP-Histone), and 3D genome organization (contact maps). AlphaGenome’s recommended variant scorers were used for each output type. For each variant and output type, the maximum absolute value was taken across all tracks within that output type as recommended by the original study authors.^8^ After removing variants with missing values, there were eight scores available (one per output type) for 8,560,287 variants, which were subsequently log-transformed and standardized (mean centered and scaled to unit variance) prior to downstream analysis.
- **Borzoi**: We incorporated 102 variant scores derived from the Sniff pipeline.^9^ Briefly, principal component analysis was applied to Borzoi’s 7,611 dimensional variant effect predictions within six functional categories (ChIP, ATAC, DNase, CAGE, RNA (ENCODE), and RNA (GTEx)) to reduce dimensionality.^9^ These 102 principal components (PCs) were used in the analysis. To handle outliers, we applied a log-modulus transformation to the PCs, sign(*x*) × log(|*x*| + 1), and then standardized the scores. In total, scores for 8,940,612 variants were available, and the full dataset can be downloaded at https://storage.googleapis.com/seqnn-share/sniff/borzoi_102_annotation_set.
- **GPN-Star**: We extracted constraint scores from GPN-Star, a genomic language model that predicts functional constraint by leveraging vertebrate, mammalian, and primate whole-genome alignments. There were 9,988,326 scores for each of the three alignments, which were subsequently encoded so that the top 0.1% most constrained values among our analysis SNPs were assigned 1 and the rest 0 (following the same procedure as the original study^6^). The data are publicly available to download at https://huggingface.co/datasets/songlab/ldsc.
- **Zoonomia**: We included two measures of genomic constraint computed on primate and mammalian alignments from the Zoonomia project.^24^ There were 9,997,231 variant scores available to download in the baselineLD (v2.3) model at https://zenodo.org/records/8292725/files/baselineLD_v2.3.zip.
- **AVI**: We evaluated the AlphaGenome variant impact score (AVI) from the AlphaGenome Atlas,^19^ which combines AlphaGenome and AlphaMissense scores with conservation and other scores to separate rare from common variants. The scores are available to download at https://deepmind.google.com/science/alphagenome.

After removing variants with missing scores in any of the annotations, the final dataset contained 7,674,000 variants. We used the PolyFun software^11^ (https://github.com/omerwe/polyfun) to calculate the corresponding LD scores for each annotation.

#### Analysis of GWAS data

To evaluate the functional annotations for common variant discovery, we analyzed summary-level data for 27 traits from the UK Biobank. We restricted our analysis to a set of well-powered and approximately independent traits (pairwise phenotypic *r*^2^ < 0.1), following the analysis described previously.^13^ These traits cover a diverse space of trait categories while also ensuring that our evaluation does not overcount the number of discoveries due to pleiotropy. GWAS summary statistics were calculated using BOLT-LMM (v2.3)^25^ applied to up to 459,324 samples of European ancestry. Additional details on quality control and preprocessing are described elsewhere.^13^ After we combined the GWAS summary statistics with our functional annotation data (MAF > 0.01) and removed variants with imputation INFO < 0.8 and the MHC region (24–45 Mb), an average of 7,061,991 SNPs remained for downstream analysis across traits.

Since sfFDR requires a set of approximately LD-independent SNPs for training, we performed LD pruning using PLINK (v2.0).^26^ Using a window size of 200 kb and an *r*^2^ threshold of 0.3 in unrelated UK Biobank participants (data field 22020), there were 616,221 LD-independent SNPs identified. A SNP was considered statistically significant if its functional *p*-value fell below the genome-wide significance threshold (*p*_f_ < 5 × 10^−8^). To conservatively define the number of independent loci, we grouped significant SNPs into clusters (*r*^2^ > 0.01 within 5 Mb) using the 1000 Genomes Phase 3 EUR reference panel, where signals that were within 100 kb of one another were collapsed to account for allelic heterogeneity. We then matched the sfFDR and standard analysis loci one-to-one by their lead SNPs (*r*^2^ > 0.01 within 5 Mb) and defined a locus as a distinct sfFDR discovery if its lead SNP was not matched to any lead SNP in the standard analysis, and vice versa.

To assess the replicability of distinct discoveries identified by sfFDR, we evaluated 11 of the 27 traits (Table S1) that had independent validation data available in the FinnGen (R12) study.^27^ A discovery was considered replicated if its FinnGen *p*-value was below the significance threshold 0.05.

## Data availability

Annotations for the analyses in this paper can be downloaded at https://doi.org/10.5281/zenodo.23063752. The data to reproduce the figures are available at https://doi.org/10.5281/zenodo.23110145. UK Biobank summary statistics are available to download at https://console.cloud.google.com/storage/browser/broad-alkesgroup-public-requester-pays/UKBB. FinnGen R12 summary statistics are publicly available at https://www.finngen.fi/en/access_results. The baselineLD and Zoonomia annotations are available at https://zenodo.org/records/8292725. Borzoi scores are available at https://storage.googleapis.com/seqnn-share/sniff/borzoi_102_annotation_set. The API to extract AlphaGenome scores is available at https://www.alphagenomedocs.com/. AVI scores are available at https://deepmind.google.com/science/alphagenome. GPN-Star annotations are available at https://huggingface.co/datasets/songlab/ldsc.

## Code availability

sfFDR is implemented in the R package sffdr (v1.1.2) and available at github.com/ajbass/sffdr. Analysis code to reproduce all results is available at github.com/ajbass/gfm_manuscript.

## Acknowledgements

This research has been conducted using the UK Biobank Resource under Application Number 98032. We want to acknowledge the participants and investigators of the FinnGen study.

## Supplementary materials

**Table S1:** The replication rate (*p* < 0.05) and total number of distinct discoveries by sfFDR with baselineLD annotations and the standard analysis in the FinnGen study (95% Wilson confidence intervals).

| Trait | sfFDR<br>(95% CI) | Distinct loci<br>(sfFDR) | Standard<br>(95% CI) | Distinct loci<br>(standard) |
| --- | --- | --- | --- | --- |
| White Blood Cell Count | 0.79 (0.69–0.87) | 82 | 0.77 (0.60–0.89) | 31 |
| Type 2 Diabetes | 0.83 (0.55–0.95) | 12 | 0.80 (0.38–0.96) | 5 |
| Red Blood Cell (RBC) Count | 0.67 (0.57–0.76) | 92 | 0.72 (0.56–0.83) | 39 |
| RBC Distribution Width | 0.51 (0.37–0.64) | 49 | 0.56 (0.34–0.75) | 18 |
| Mean Corpuscular Hemoglobin | 0.73 (0.60–0.82) | 62 | 0.73 (0.54–0.86) | 26 |
| Hypothyroidism | 0.96 (0.79–0.99) | 23 | 0.80 (0.38–0.96) | 5 |
| Height | 0.64 (0.55–0.71) | 129 | 0.50 (0.40–0.60) | 96 |
| Eosinophil Count | 0.47 (0.34–0.60) | 51 | 0.50 (0.34–0.66) | 32 |
| Cardiovascular | 0.51 (0.36–0.66) | 41 | 0.53 (0.30–0.75) | 15 |
| BMI | 0.83 (0.75–0.89) | 101 | 0.62 (0.51–0.72) | 74 |
| Autoimmune Disease | 0.87 (0.62–0.96) | 15 | 1.00 (0.61–1.00) | 6 |
| All traits | 0.69 (0.65–0.72) | 657 | 0.61 (0.56–0.66) | 347 |

**Figure S1:**
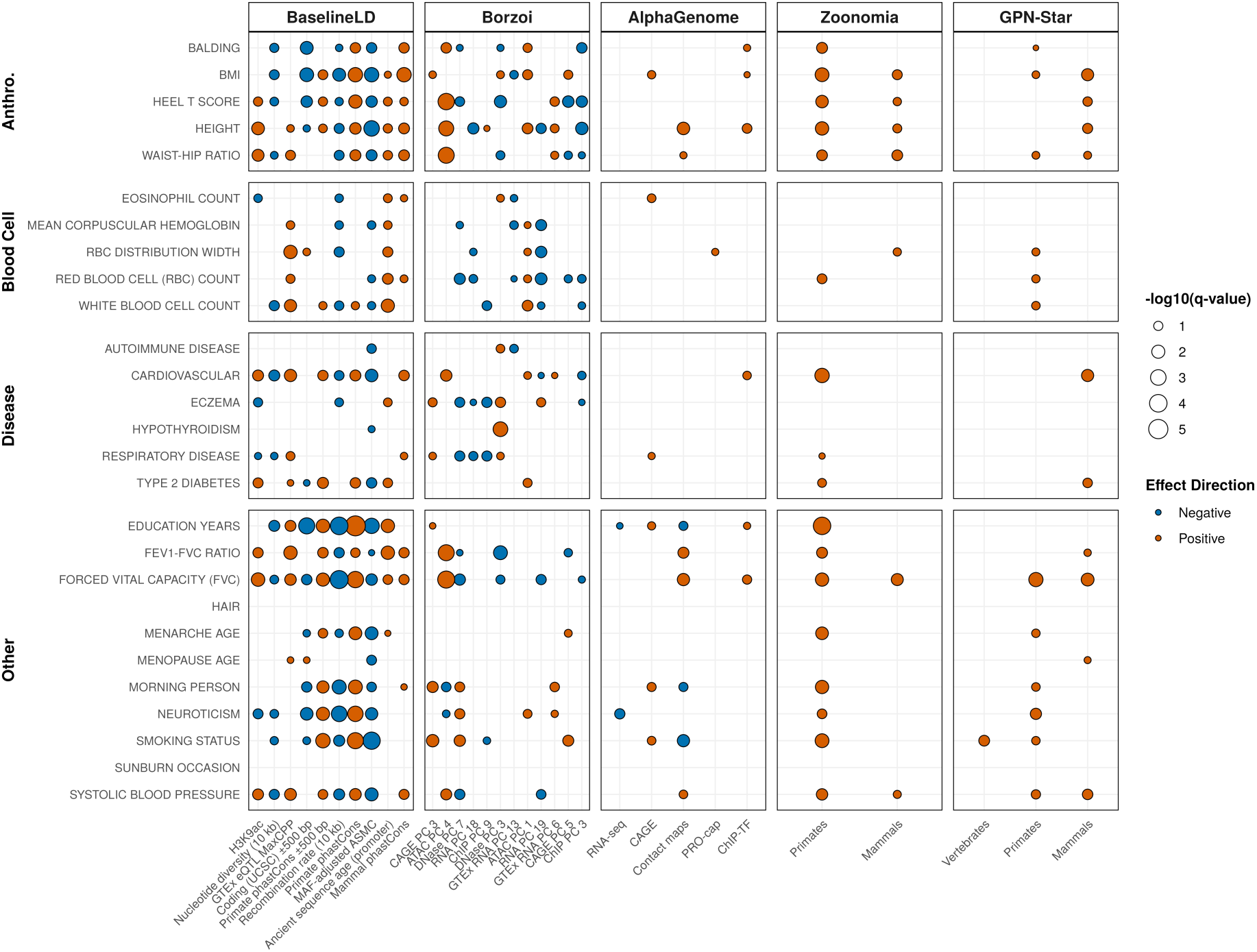
Key drivers of functional enrichment across the 27 UK Biobank traits. Statistically significant explanatory variables (FDR < 0.2) identified via ridge regression to estimate the expected *χ*^2^ across traits (rows) and annotation models (columns). Bubble size is scaled by −log10(*q*-value), and fill color denotes the sign of the coefficient. For baselineLD annotations, the top 10 significant predictors across multiple classes were selected. For Borzoi, only the top two principal components per modality were selected. Note that baselineLD annotations are included in the models for Borzoi, AlphaGenome, Zoonomia, and GPN-Star.

**Figure S2:**
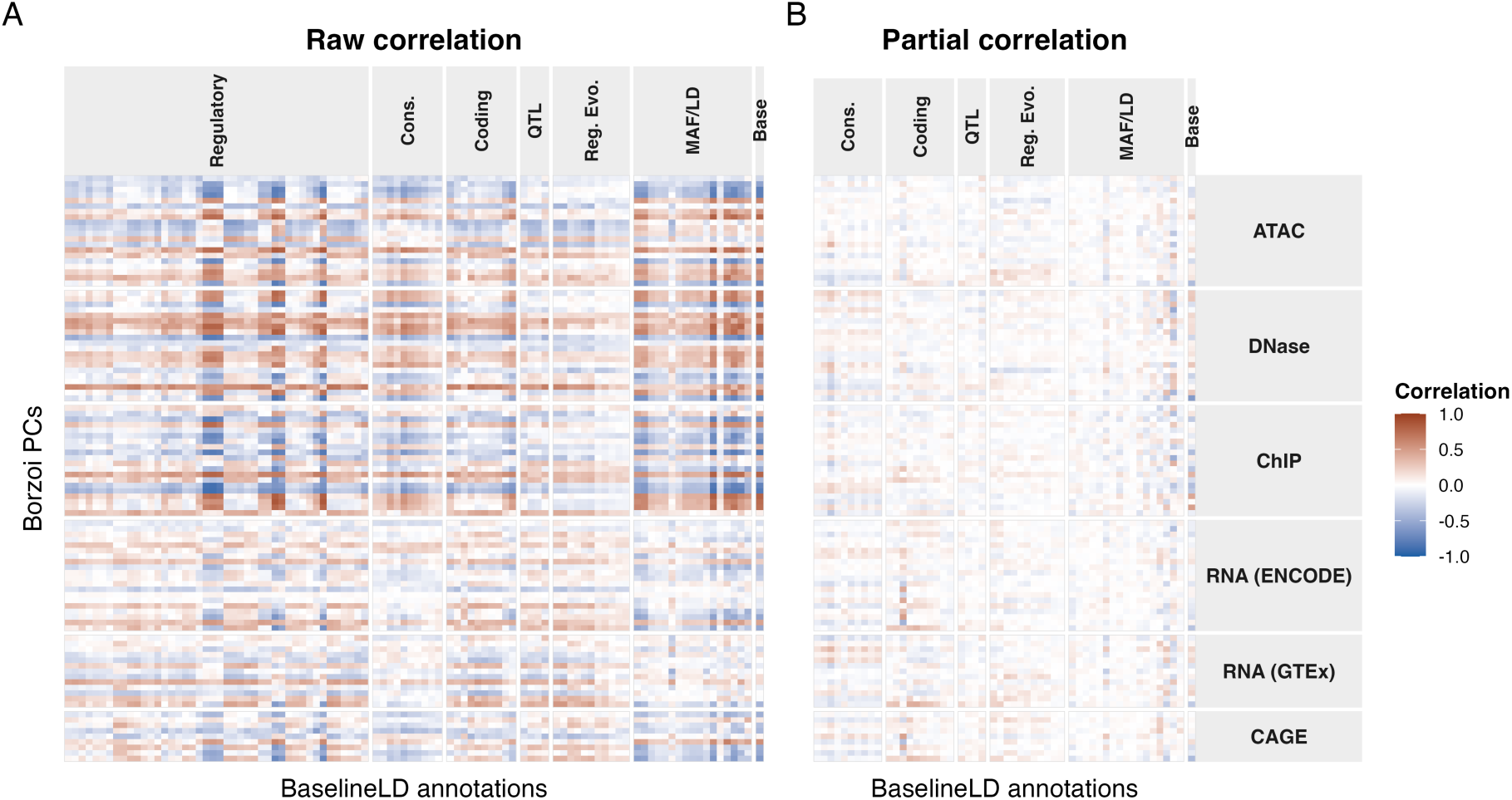
Borzoi PCs substantially overlap with baselineLD regulatory annotations. Pearson correlations between LD scores of Borzoi PCs (rows) and baselineLD annotations (columns). (A) Raw correlations between annotations. (B) Partial correlations between Borzoi’s PCs and baselineLD annotations conditioning on the regulatory annotations. Color denotes the correlation coefficient on a common scale (blue, negative; red, positive).

**Figure S3:**
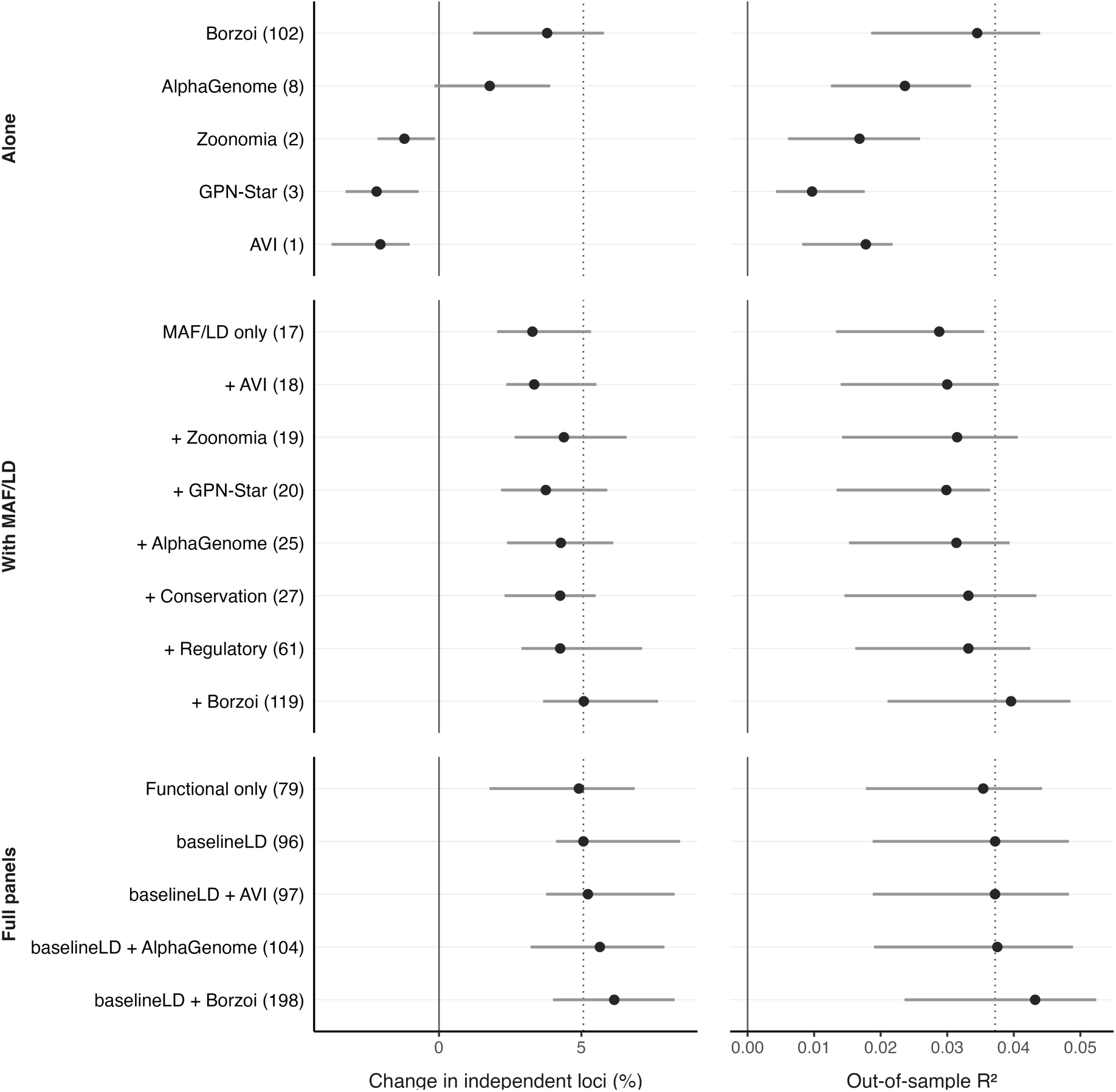
Evaluating functional annotations beyond genetic architecture in 27 UK Biobank traits. The change in the number of independent loci relative to standard *p*-values (left) and out-of-sample *R*^2^ (right). The points are the medians and bars are interquartile ranges. The solid line denotes no change while the dotted line denotes baselineLD. Note that parentheses give the number of annotations in each set and the functional category removes the MAF and LD-architecture annotations from baselineLD. AVI denotes the standardized AlphaGenome variant impact score.^19^

## References

1. Consens, M. E. et al. Transformers and genome language models. Nat. Mach. Intell. 7, 346–362 (2025).

2. Benegas, G., Ye, C., Albors, C., Li, J. C. & Song, Y. S. Genomic language models: opportunities and challenges. Trends Genet. 41, 286–302 (2025).

3. Jumper, J. et al. Highly accurate protein structure prediction with AlphaFold. Nature 596, 583–589 (2021).

4. Cheng, J. et al. Accurate proteome-wide missense variant effect prediction with AlphaMissense. Science 381, eadg7492 (2023).

5. Christmas, M. J. et al. Evolutionary constraint and innovation across hundreds of placental mammals. Science 380, eabn3943 (2023).

6. Ye, C. et al. Predicting genome-wide functional constraints with GPN-Star. Nature 10.1038/s41586-026-11005-5 (2026).

7. Linder, J., Srivastava, D., Yuan, H., Agarwal, V. & Kelley, D. R. Predicting RNA-seq coverage from DNA sequence as a unifying model of gene regulation. Nat. Genet. 57, 949–961 (2025).

8. Avsec, Ž., et al. Advancing regulatory variant effect prediction with AlphaGenome. Nature 649, 1206–1218 (2026).

9. Srivastava, D. et al. Borzoi-informed fine mapping improves causal variant prioritization in complex trait GWAS. Preprint at bioRxiv 10.1101/2025.07.09.663936 (2025).

10. Li, X. et al. Dynamic incorporation of multiple in silico functional annotations empowers rare variant association analysis of large whole-genome sequencing studies at scale. Nat. Genet. 52, 969–983 (2020).

11. Weissbrod, O. et al. Functionally informed fine-mapping and polygenic localization of complex trait heritability. Nat. Genet. 52, 1355–1363 (2020).

12. Dey, K. K. et al. Evaluating the informativeness of deep learning annotations for human complex diseases. Nat. Commun. 11, 4703 (2020).

13. Kichaev, G. et al. Leveraging polygenic functional enrichment to improve GWAS power. Am. J. Hum. Genet. 104, 65–75 (2019).

14. Bass, A. J. & Wallace, C. Exploiting pleiotropy to enhance variant discovery with functional false discovery rates. Nat. Comput. Sci. 5, 769–781 (2025).

15. Gazal, S. et al. Linkage disequilibrium dependent architecture of human complex traits shows action of negative selection. Nat. Genet. 49, 1421–1427 (2017).

16. Finucane, H. K. et al. Partitioning heritability by functional annotation using genome-wide association summary statistics. Nat. Genet. 47, 1228–1235 (2015).

17. Marderstein, A. R. et al. Decoding common and rare noncoding variant effects across cellular and developmental contexts. Nat. Genet. 58, 1517–1529 (2026).

18. Drusinsky, S., Whalen, S. & Pollard, K. S. Deep-learning prediction of gene expression from personal genomes. Genome Biol. 27, 19 (2026).

19. Cheng, J. et al. AlphaGenome Atlas: in silico mutagenesis of the entire human genome improves prioritization and interpretation of non-coding variants. Preprint at medRxiv 10.64898/2026.09.16.26363192 (2026).

20. Hu, E. J., et al. LoRA: low-rank adaptation of large language models. Preprint at arXiv 10.48550/arXiv.2106.09685 (2021).

21. Sirugo, G., Williams, S. M. & Tishkoff, S. A. The missing diversity in human genetic studies. Cell 177, 26–31 (2019).

22. Martin, A. R. et al. Current clinical use of polygenic scores will risk exacerbating health disparities. Nat. Genet. 51, 584–591 (2019).

23. Chen, X., Robinson, D. G. & Storey, J. D. The functional false discovery rate with applications to genomics. Biostatistics 22, 68–81 (2021).

24. Sullivan, P. F. et al. Leveraging base-pair mammalian constraint to understand genetic variation and human disease. Science 380, eabn2937 (2023).

25. Loh, P.-R., Kichaev, G., Gazal, S., Schoech, A. P. & Price, A. L. Mixed-model association for biobank-scale datasets. Nat. Genet. 50, 906–908 (2018).

26. Purcell, S. et al. PLINK: a tool set for whole-genome association and population-based linkage analyses. Am. J. Hum. Genet. 81, 559–575 (2007).

27. Kurki, M. I. et al. FinnGen provides genetic insights from a well-phenotyped isolated population. Nature 613, 508–518 (2023).

